# Engineered Microbes to Sense and Respond to Enterotoxigenic Escherichia coli

**DOI:** 10.1101/348573

**Authors:** A. Bete, J. Carter, C. Davis, J. Dong, M. Herrmann, H. Jesse, D. McDonald, P. Menart, A. Poole, A. Smith

## Abstract

Every year, Enterotoxigenic *Escherichia coli* (ETEC), the most common form of traveler’s diarrhea, affects thousands of military personnel deployed overseas. The goal of this research was to engineer non-pathogenic *E. coli* to sense ETEC, respond to its presence, and package the non-pathogenic *E. coli* in a cellulose matrix to enable environmental detection of ETEC. Two plasmids were created: ‘sense-respond’; and ‘packaging’. The sense-respond plasmid detected autoinducer 2 (AI-2), a quorum sensing molecule created by most ETEC strains, by expressing LsrR which switches on the Lsr promoter. Activation of the Lsr promoter expresses superfolder green fluorescent protein (sfGFP), indicating the presence of ETEC. The packaging plasmid expresses a fusion protein consisting of curli fibers and cellulose binding domains. These modified surface proteins permit the bacteria to bind to cellulose, encapsulating the sense-response module. This genetically engineered machine could be deployed in both the internal and external environment to detect ETEC.

## Introduction

Many travelers suffer from diarrhea when traveling overseas because of contact with foreign microbes. Microbes that are known to cause diarrhea include Enterotoxigenic *E. coli*, Enteroaggregative *E. coli*, Campylobacter, Norovirus, Rotavirus, and some unidentified bacteria. Exposure to these microbes can result in deployed personnel and civilian travelers becoming ill and experiencing various symptoms, such as severe diarrhea, fevers, stomach cramps, and vomiting. Approximately, 60% of deployed personnel experience some form of these symptoms, commonly known as “Traveler’s Stomach” [paper in progress of publishing]. Within the 60% of deployed personnel, 47% experience three or more episodes. Consequently, deployed personnel suffer from operational impacts such as reduced job performance, restriction to bed rest, hospitalization, missed patrols, or requiring backup personnel. Enterotoxigenic *E. coli* (ETEC) is the leading known cause of Traveler’s Stomach for both deployed military personnel and civilian travelers. ETEC is the most common toxic form of Traveler’s Stomach for deployed personnel and produces two toxins, heat labile and heat stable [5].

Nearly 76% of travelers tested ETEC-positive in their stool samples after visiting three foreign countries (India, Guatemala, and Mexico). The locations of different developing countries determine the intensity of the effect of ETEC and its toxins. Studies have shown where Enterotoxigenic *E. coli* is present the most and how the effects of ETEC change from country to country. As countries continue to develop, ETEC remains present in communities, affecting many civilians in the community that are not native to the area. [5].

Removing genes from one plasmid and inserting them into another resembles a form of plug and play. Many variations of genes exist, and biologists can combine genes in a multitude of ways. For example, green fluorescent proteins (GFPs) are commonly used to create a response that can be viewed optically. There are several different types of GFPs, such as super folder GFP and GFPa1. To view the GFP, scientists place the samples under ultraviolet light. Fluorescent proteins displaying these colors permit the viewer to better differentiate between the green tint of a bacterial colony and the fluorescing protein [1].

Quorum sensing refers to a way that bacteria communicate with each other using different types of signals. Quorum sensing molecules can be specific to bacteria populations and are tuned to various concentrations. The concentration of these molecules corresponds to a high density, or quorum. When the concentration reaches a certain quantity, the whole population initiates a coordinated expression of specific genes. Autoinducer-2 is a quorum sensing molecule that is produced by different kinds of *E. coli* including Enterotoxigenic *E. coli* [8].

Encapsulation of a cell is made possible through extracellular binding domains that can bind to specified materials. An ideal material that is widely studied and characterized is celluose. It is not broken down within the GI tract and is produced naturally, making it a viable means of encapsulation. Several types of cellulose specific binding domains exist. A double cellulose binding domain (dCBD) is two cellulose binding domains that have been combined. There are two linkers and two binding domains. The initial part is the linker which allows the double cellulose binding domain to connect to different fibers or proteins. This linker is essential to the functioning of the cellulose binding domain (CBD) and facilitates reactions and influences the effective concentrations of reactants (Linder). The second component is the binding domain, which acts like a strip of velcro. This allows for twice as much binding to the cellulose. Having twice as many binding domains adds to the possibility of the bacterium remaining connected to the cellulose matrix and not being released into the body or into the environment. Such a characteristic is important because of the public concern regarding the safety of genetically modified organisms (GMOs).

A series of curli fibers form a mesh around a cell and displays binding sites along the entirety of the fiber, allowing the cell to attach to certain surfaces. These fibrous proteins are considered the first workable amyloid discovered. Curli fibers are associated with organisms that use fibers to attach to the host (9). Curli fibers are the product of separate subunits called curli specific genes (*csg*) with a corresponding letter for each subunit. Two subunits of curli actually protrude from the bacterium, the first being the csgA subunit. The *csg*A constitutes most of the curli fiber; it acts as the building blocks of the fiber. According to Chapman Laboratory, csgA is composed of five layers of fibers that are connected and twisted, similar to a spring. Although few curli fibers protrude from the bacterium, the number of *csg*A subunits that are produced, one on top of the other, form extensive strands of curli. The bacterium will have a fibrous mesh surrounding almost the entire outer layer. CsgA begins assembling on the initial block of *csg*B. CsgD is a positive transcriptional regulator of *csg*A and *csg*B. The purpose of *csg*C is unknown. The last three subunits: *csg*E, *csg*F, and *csg*G are subunits about which information is still being gathered. CsgG was recently discovered to assist in the secretion and stability of *csg*A and *csg*B [2]. Curli fibers are incredibly robust; for example, they can withstand boiling in detergents and incubation in solvents. This makes them extremely versatile and able to work in many types of environments. These fibers are 10-40% of the total biovolume in a biofilm, which means a significant amount of a biofilm could be engineered. The curli is the most widely studied and a straightforward platform to work with for *E. coli* [7].

The csgA subunit was found to be a viable way to attach the dCBD domains to curli fibers to ensure that the bacterium is encapsulated in cellulose. As there are thousands of *csg*As that are built up on top of each other, this would allow for the dCBD to be much more abundant because it will be able to connect its linker to the ends of each of the five fibers that make up a single *csg*A subunit. Therefore, each *csg*A unit will contain five dCBDs or ten CBDs in total on each *csg*A subunit[2].

CsgA have the ability to attach many dCBD to each fiber which strengthen the connection to the cellulose matrix. This would permit the module inside the cellulose pellicle to pass through the human GI tract unharmed. Having the module encased in such a pellicle should reassure the person ingesting the GMO that the module would remain encased in cellulose throughout their digestive tract. Furthermore, the cellulose pellicle would allow the testing of water conditions. The module could be placed into a container of water in order to detect pathogenic strains of bacteria, indicating whether the water is safe to drink. Throughout this process the module would remain enclosed in the cellulose pellicle [4].

We propose engineering a probiotic strain of *E. coli* for use in diagnostic testing. This engineered *E. coli* will produce GFP in response to the presence of ETEC and will be encapsulated in cellulose.

## Materials and Methods

### Cell Strains/Chassis

*Escherichia coli* DH5-alpha and TOP10 competent cells were primarily used as the chassis for plasmid cloning. These were chosen due to their growth rate, availability, and high plasmid retention. The plasmids created were optimized for *E. coli*.

### Plasmid Cloning

Plasmid components were initially obtained from various Air Force collaborators. The components for the *csg*A gene was derived from *E. coli* Nissle 1917 genomic DNA, and the double cellulose binding domain and super folder GFP was taken from the iGEM parts registry. The three created components were then sequenced and ordered as gBlocks from IDT and inserted into the pSB1C3 via ligation.

Plasmid cloning took place using New England Biolabs cut enzymes and T4 DNA Ligation kits. After DNA was enzymatically cut, the sample was processed on an agarose gel (1% TAE buffer) and the desired DNA was cut and processed using a Qiagen Gel Extraction kit. Newly created plasmids were then isolated from whole cells using Qiagen QIAprep Spin Miniprep kits. Plasmids were then transformed into competent cells purchased from New England Biolabs. Cells were incubated on ice for 20 minutes, heat shocked at 42 C for 45 seconds, then placed on ice for two minutes. SOC media was added for the recovery phase and the cells were incubated at 37 C (at 215 rpm) for 2 hours, then plated.

### Fluorescence Assays

DNA samples were quantified for quality and quantity using a Nanodrop spectrophotometer. Optical density and fluorescence of cell samples were taken using a SpectraMax M5 Plate Reader. The optical density wavelength was set at 600 nm and blanked with plain LB-broth.

### GFP Project

The fluorescent output of GFPa1 and sfGFP were compared to each other and measured using the plate reader method described above. Samples were inoculated at time 0 with an inducer (3OC12) and then measured on a plate reader at 2, 4, 6,8, and 24 hour time points.

### Growing Cellulose

Cellulose was grown from *Gluconacetobacter hansenii.* The bacteria was grown inside well plates containing agar. Half of the plates were incubating while stationary and the other half were incubating while oscillating at 27 C and 200 rpm. The cellulose grown was then extracted and put into 50ml centrifuge tubes. The tubes were lyophilized in order to dry the cellulose into pellicles over night.

## Results

### Engineered Escherichia coli to Detect 3OC12 and Respond with Superfolder Green Fluorescent Protein (BioBrick Part: BBa_K2522001)

A test plasmid was created to examine the concept of a plasmid construct using commonly and readily available materials due to the lack of known facts related to ETEC. The test plasmid contained a PLas promoter activated by quorum sensing molecule 3OC12. Once the PLas promoter was activated it produced green fluorescent protein A1 [11]. Figure 1 demonstrates the 3OC12 activating the PLas promoter, allowing the RBS to initiate the production of GFPa1. Figure 2 shows the working construct developed to produce green fluorescent protein A(GFPa1) in the presence of the quorum sensing molecule 3OC12. Through both experimentation and sequencing, the DNA of the working construct was confirmed to work as theoretically planned. Image 1 displays the experimental data. Lab results illustrate that GFPa1 was only expressed in the presence of 3OC12, which shows the specificity of the PLas promoter. Dimethyl sulfoxide (DMSO), a solvent that dissolves 3OC12, alone did not produce a response from the PLas promoter [12].

**Figure 1.**
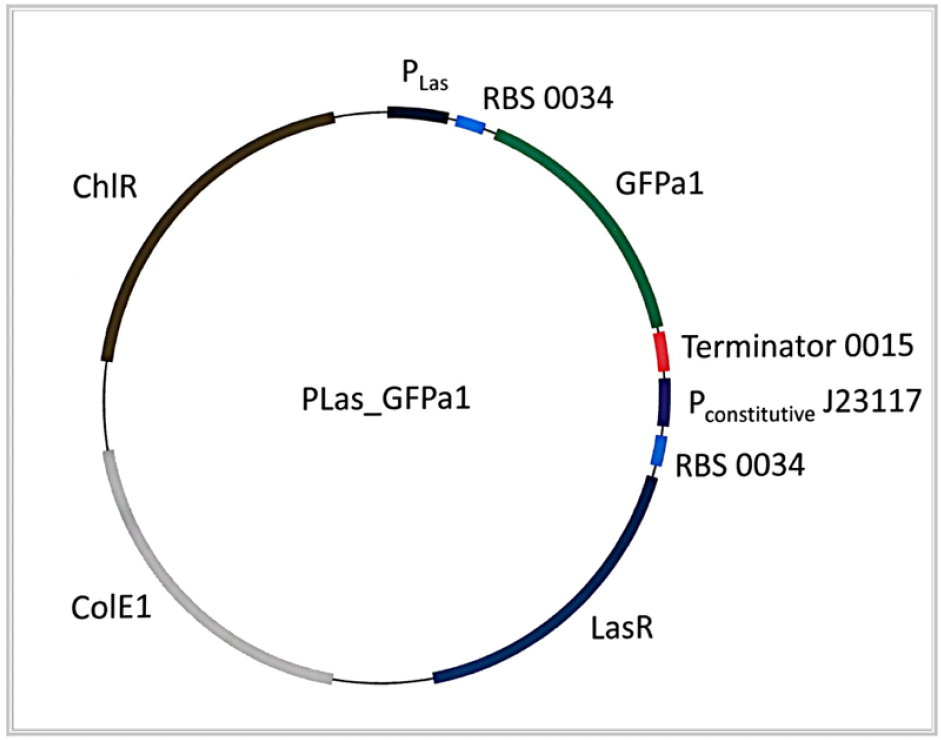
Plasmid Map: The plasmid contains a PLas promoter, ribsomal binding site, GFPa1, a terminator, followed by a constitutive promoter to continually produce LasR.

**Figure 2.**
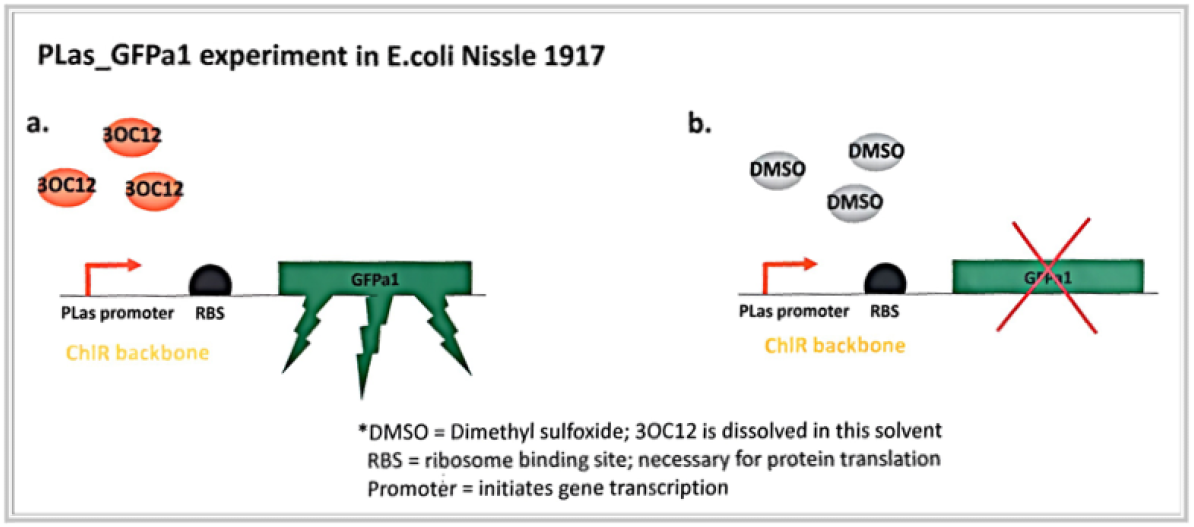
The left construct demonstrates the PLas promoter as specific to 3OC12. In the right construct the PLAs promoter is not activated in the presence of the control solvent DMSO.

**Figure 3a.**
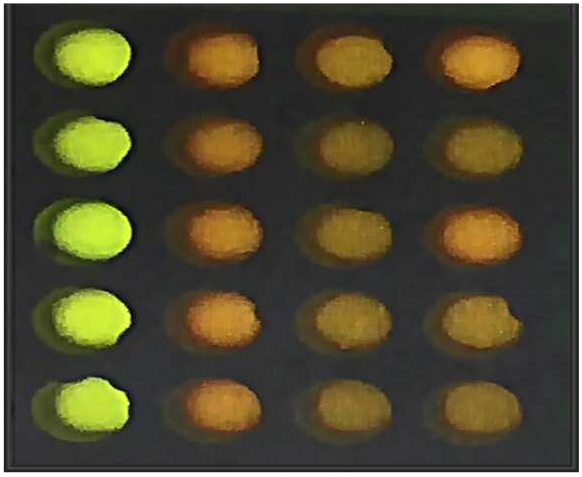
The plate reader results of the induced PLas-GFPa1 Plasmid. The columns in order from left to right: GFPa1 Colony + 3OC12, GFPa1 Colony + DMSO, LB + 3OC12, LB + DMSO.

**Figure 3b.**
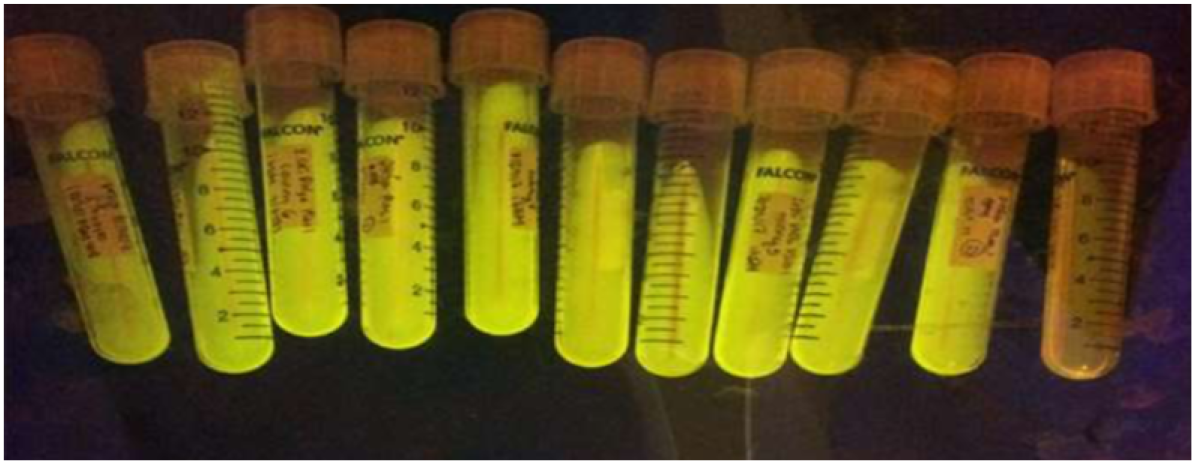
The picture shows the bright fluorescence production of the GFPa1.

After the successful production and testing of the PLas-GFPa1, the production rate of the fluorescence in this plasmid was not applicable to the needs of the final plasmid. The final plasmid would need to have a faster reaction time for the diagnostic environment for our future plasmid. In order to improve the fluorescent aspect of the plasmid, it was determined to do comparative study testing the reaction rates of GFPa1 and sfGFP, a green fluorescent protein suspected to have a faster rate of fluorescence [11].

### Comparison Study of Green Fluorescent Protein A1 and Superfolder Green Fluorescent

**Figure 4a, 4b.**
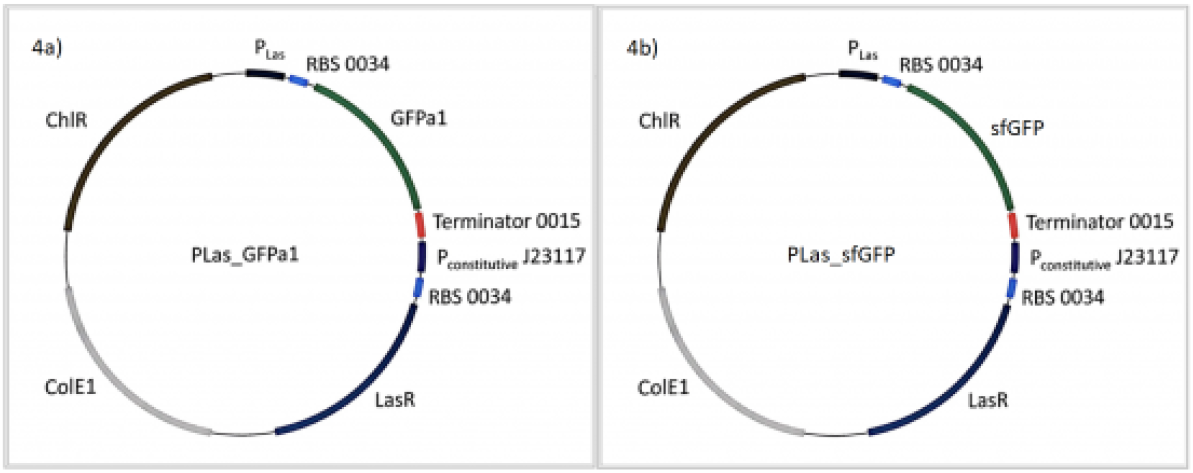
The left plasmid map (4a) is the PLas promoter which begins the production of the GFPa1 in the presence of 3OC12. The right plasmid map (4b) is the PLas Promoter which begins the production of sfGFP.

The comparison study was performed using the same plasmid except with sfGFP in place of the GFPa1. Figure 4a shows the original test plasmid and Figure 4b shows the adapted plasmid to be used in the comparison study. The study was conducted using a plate reader; in addition, fluorescence was read every hour for 8 hours and then a final reading was taken at 24 hours. Figure 5 exhibits the fluorescence test tabularly. In examining the data the results supported the hypothesis: the sfGFP demonstrated a significantly faster reaction time with the fluorescence rate increasing at Hour 4. The GFPa1 did not produce a high fluorescence rate until Hour 8. The final reading also indicated sfGFP had a much higher end reading at Hour 24 then the GFPa1. These results suggested that sfGFP would be a more applicable form of GFP than GFPa1. The final plasmid to be sequenced and submitted was the PLas-sfGFP. (Sequencing
results: http://parts.igem.org/Part:BBa_K2522001)Once the test plasmid/construct was created, research was 2 performed to find a quorum sensing molecule specific to Enterotoxigenic *Escherichia Coli* (ETEC) [8].

**Figure 5.**
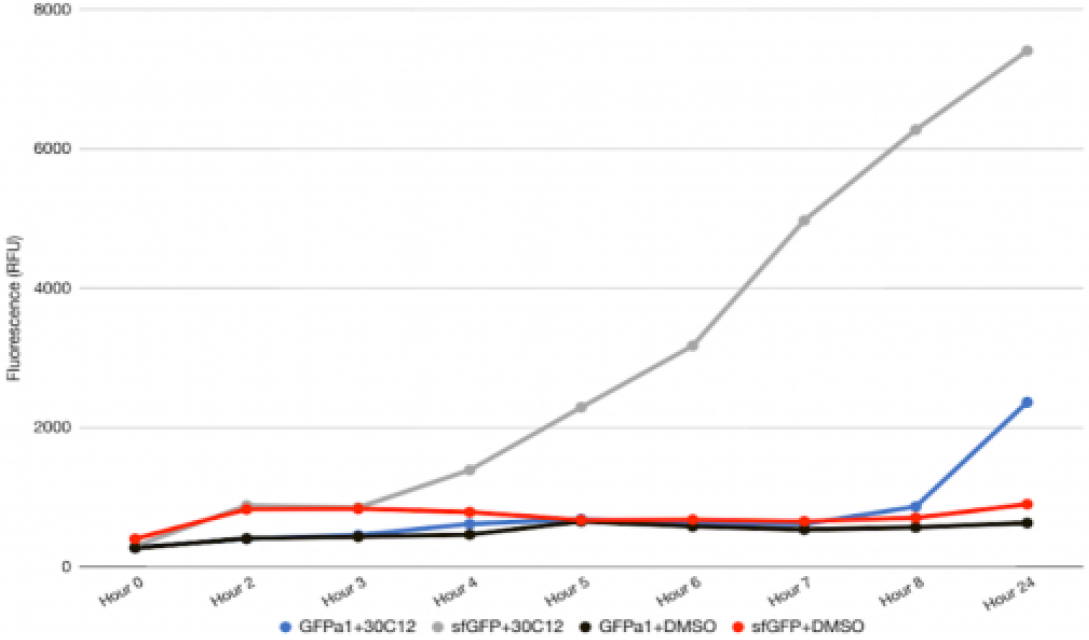
The superfolder green fluorescent protein leads to a faster reaction at hour 4. The 3OC12 was added to begin the production of the GFPa1 and sfGFP. The DMSO was in added in the controls

### Engineered *Escherichia coli* to Detect AI-2 and Respond with Superfolder Green Fluorescent Protein (BioBrick Part: BBa_K2522002)

An iGEM Part Registry search located a part containing a PLsrA promoter and a yellow fluorescent protein response, BioBrick Part Number: BBa_K117008. The PLsr promoter is activated by autoinducer 2, a quorum sensing molecule for several pathogenic gut diseases including ETEC (8). The part was then adapted for our application by performing a digestion and ligation to replace the YFP for sfGFP. Figure 6 displays a plasmid map comparison of the iGEM Part and the adapted plasmid containing the sfGFP. The plasmid was grown up overnight in supernatant from an E.coli Nissle 1917 sample, which in theory should contain AI-2. However, the test was inconclusive due to contamination of bacterial growth in pure LB broth used as a blank. While confirmed to match the theoretical design via sequencing, only one experiment was performed with the plasmid due to time constraints. (Sequencing results: http://parts.igem.org/wiki/index.php?title=Part:BBa_K2522002) To contain *E. coli* within a cellulose matrix, a plasmid was developed using the curli-specific-gene A (*csg*A) extracted from the genomic DNA of *E. coli* Nissle (Figure 8) and a double cellulose binding domain gene taken from the iGEM DNA Distribution Kit. We designed the plasmid to enable an *E. coli* bacterium to produce curli fibers with a double cellulose binding domain attached to each *csg*A subunit so that the bacterium can bind itself to cellulose. To obtain the desired plasmid expressing the fusion protein of curli fibers with double cellulose binding domain, the double cellulose binding domain (dCBD) was extracted from the iGEM 2017 parts kit. After the extraction, the dCBD gene was fused with the curli fiber gene (cgsA). This was accomplished by ligating the two genes and removing the stop codon of the *csg*A gene. As a result, the *csg*A gene would flow continuously into the dCBD gene to form curli fibers that bind to cellulose (Figure 8).

**Figure 6.**
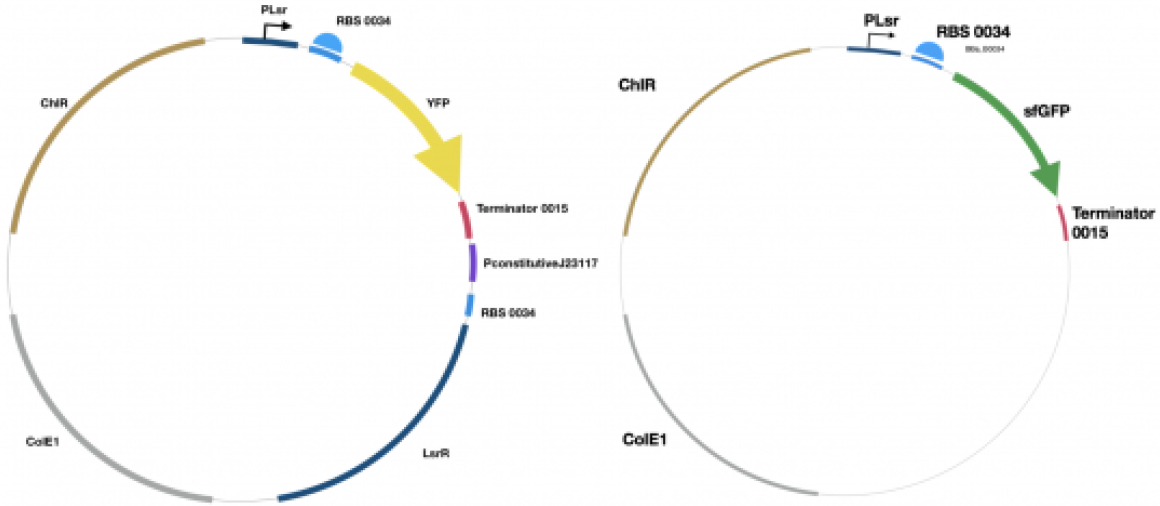
Plasmid Map. The left plasmid map is the original iGEM part with a response of YFP. The right plasmid map is the PLsr promoter begins the production of the sfGFP in the presence of autoinducer 2.

**Figure 7.**
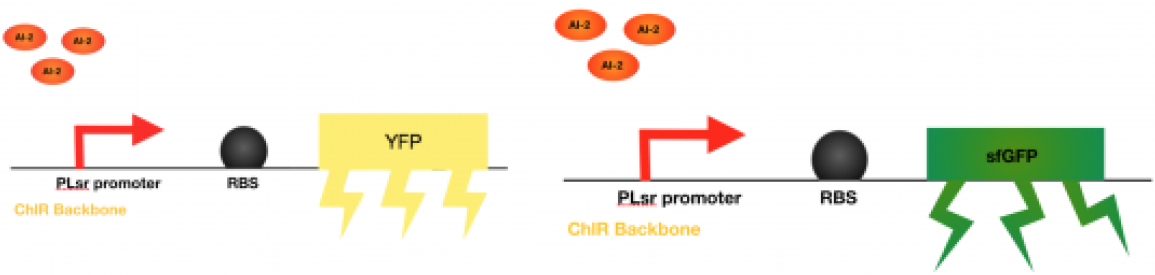
The left construct demonstrates the original iGEM part (BioBrick ID: BBa_K117008) with the PLsrA Promoter activating YFP production. The right construct demonstrates the adapted version of the PLsrA promoter activating sfGFP production.

**Figure 8.**
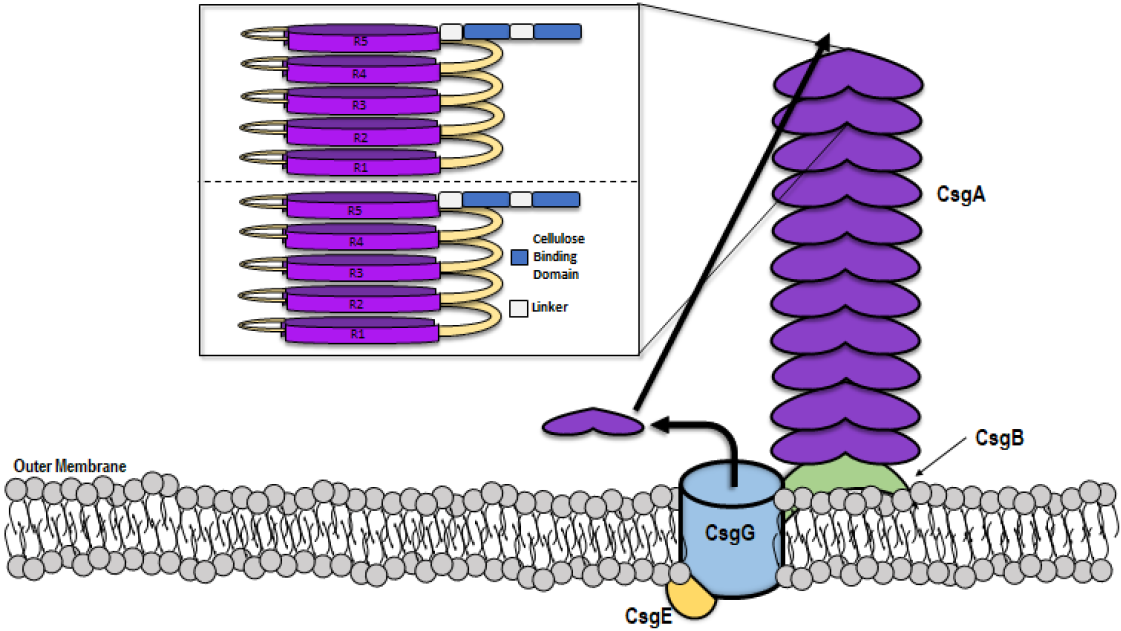
This shows the process and components of building a curli fiber. This figure is adapted from Chapman Labs. It shows the curli specific genes that are needed for the fiber. The construct of the CsgA is shown with a visual addition of the double cellulose binding domain that was found in the iGEM kit. The double cellulose binding domain is shown with the linker used for expression purposes.

The fusion gene was then cloned into a pBAD backbone and transformed into competent cells. The gene that has been created is located between the restriction sites of Nco1 and HindIII, with a Kpn1 restriction site between the *csg*A and dCBD DNA sequences. The *csg*A-dCBD was created in the pCR-Blunt II-TOPO vector from NEB Biolabs. The *csg*A-dCBD was later removed from the Blunt II- TOPO vector and cloned into the pBAD plasmid. This new plasmid was named pJAAM01, but contained extra DNA bands due to an extra restriction site that was not accounted for when restriction digest was performed with the pCR-Blunt II-TOPO plasmid. As the end of internship was approaching, a *csg*A-dCBD gBlock was synthesized and ordered through IDT, Inc. The *csg*A-dCBD gBlock was assembled into pSB1C3 plasmid backbone via Gibson DNA assembly. The original plan was to subclone the *csg*A-dCBD gene into pBAD His A vector to create the correct pJAAM01 (Figure 9). The other genes and proteins needed to create a curli fiber are accounted for in the genome of the *E. coli* and does not need to be used with the fusion gene. The correct sequence of the *csg*A-dCBD gene in pSB1C3 was submitted to the iGEM parts registry. However, no further analysis was performed on the plasmid due to time constraints of the internship ending.

**Figure 9.**
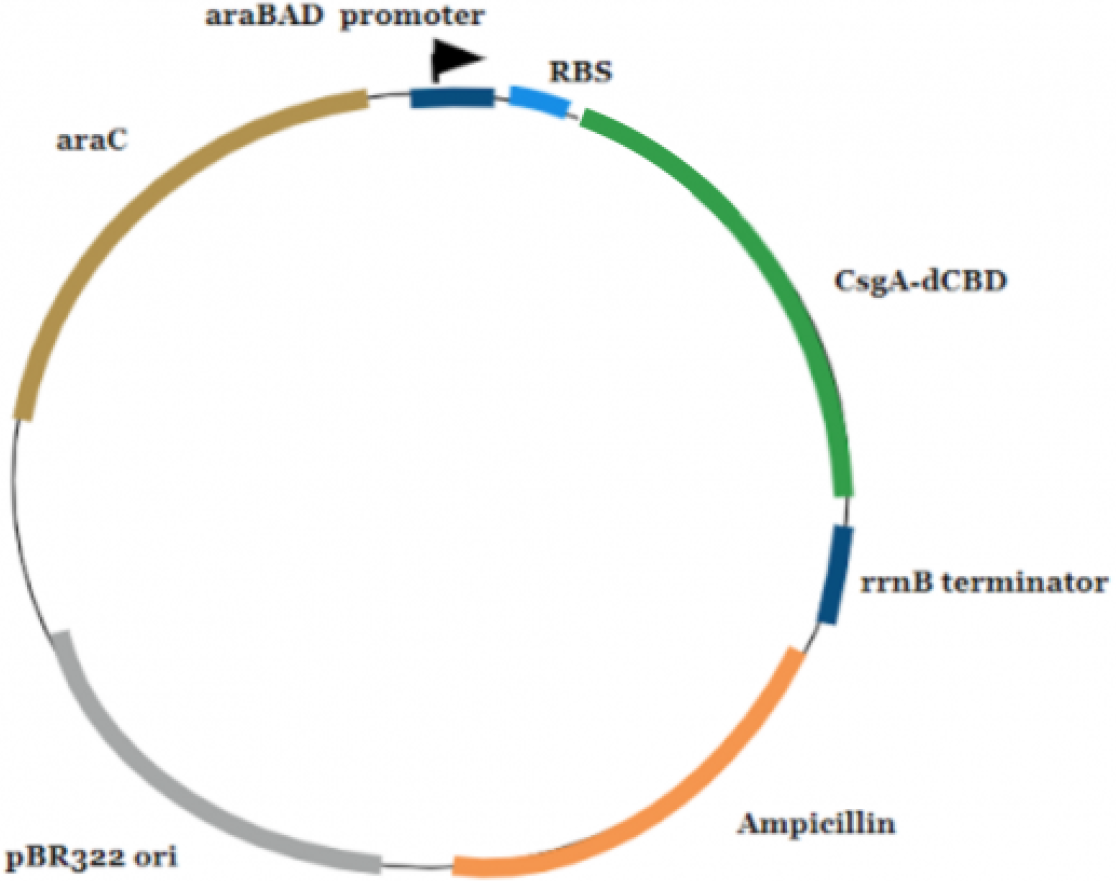
pJAAM02 Plasmid Figure 9 is the completed pJAAM02 plasmid. The backbone of this plasmid is pBAD which contains an origin site for replication purposes, the araBAD promoter, ampicillin resistance, and the inserted csgA-dCBD fusion gene. The ampicillin resistance is necessary to force the E. coli cells to replicate the plasmid. This plasmid was sent to the iGEM parts registry with the part number BBa_K2522000. This plasmid was named pBAD-csgA-dCBD before renaming it to pJAAM.

The final plasmid was named pJAAM02. This portion of the project concluded with the correct sequencing of the plasmid, pJAAM02, containing the desired fusion gene of *csg*A-dCBD seen in Figure 8. This plasmid was sent to the iGEM parts registry for distribution to all iGEM teams with the part number BBa_K2522000.

## Conclusions/Discussion

In conclusion, the US_AFRL_CarrollHS team created three plasmids. The first PLas_GFPa1 plasmid is capable of producing a response with GFPa1 in the presence of 3OC12. Next an ETEC specific plasmid was designed with the modified PLsrA_sfGFP plasmid which responds with superfolder GFP in the presence of AI-2. Both parts were sequenced and submitted to the iGEM parts database. The third plasmid was the pJAAM02 which is a pBAD backbone vector that contains the csgA-dCBD insert. This plasmid will help contain the nontoxic *E. coli* in a cellulose matrix.

This project is aimed toward preventing deployed Air Force personnel as well as civilian travelers from contracting travelers’ stomach. It can help the airmen avoid water contaminated with ETEC by placing a cellulose matrix that incorporates the sense and response module while binding to the cellulose in potentially contaminated water. Our module could also be useful in the human body. Although no human testing has been done, an airmen could ingest the cellulose capsule to determine if ETEC is present in the body. The cellulose was designed with the goal of travelling through the human GI tract. The pJAAM02 plasmid would keep the *E. coli* cells bound to the cellulose. Once the capsule travels through their GI tract, the capsule would fluoresce green in the presence of ETEC. This would then allow the airmen to get medical help before the symptoms of travellers stomach would remove them from their field work. If no ETEC is detected, the module would be released harmlessly through stool. The environmental impacts of releasing our capsule into the environment have yet to be determined, but our next goals included designing a killswitch within the plasmid.

In the future, the team will continue to further test both plasmids with the eventual goal of testing in a conditions similar to the GI tract or extreme temperature, humidity, or weather environmental settings. Various modelled promoters will also be toggled to see which one is most accurate and efficient. Additionally, the Packaging plasmid will be tested with congo red to see if the *E. coli* will produce curli fibers. Once the curli fibers are made, cellulose will be grown to bind to the cellulose matrix. Plans also include testing what conditions allow for the bacteria toescape as well as testing the strength of the binding in similar conditions to that of the body (acid levels, temperature, etc.).

## Acknowledgements

We would like to thank the United States Air Force for providing the grant that allowed us to participate in iGEM this year, as well as for providing the research labs and mentors that carried us through the project. We would like to thank our Air Force Research Lab mentors: Annastasia Bennett, Mattie Carter, Jorge Chavez, Mike Goodson, Wendy Goodson, Svetlana Harbaugh, Chia Hung, Nancy Kelley-Loughnane, Rachel Krabacher, Rajesh Naik, Vanessa Varaljay, and Doug Carter.

We would like to thank our teachers at Carroll High School: Dr. Martha Carter, Dr. Caroline Metosh-Dickey, and Dr.Christina O’Malley, along with our principal, Matt Sableski. Carroll High School provided us meeting space, allowed us to distribute surveys to students, and granted us the opportunity to practice presenting to members of the faculty.

We would also like to thank UES for sponsoring our iGEM team, permitting us to work in their meeting rooms, and letting us use their lab space for training on safety and basic techniques. We would like to thank Elizabeth Nijak and Dr.Melanie Tomczak for assisting us with logistics at UES

We would like to thank Integrated DNA Technologies (IDT) for providing free DNA sequencing for our plasmids.

